# The *Ficus erecta* genome to identify the Ceratocystis canker resistance gene for breeding programs in common fig (*F. carica*)

**DOI:** 10.1101/749788

**Authors:** Kenta Shirasawa, Hiroshi Yakushiji, Ryotaro Nishimura, Takeshige Morita, Shota Jikumaru, Hidetoshi Ikegami, Atsushi Toyoda, Hideki Hirakawa, Sachiko Isobe

## Abstract

*Ficus erecta*, a wild relative of common fig (*F. carica*), is a donor of Ceratocystis canker resistance in fig breeding programs. Interspecific hybridization followed by recurrent backcrossing is an effective method to transfer the resistance trait from wild to cultivated fig; however, this is time consuming and labor-intensive for trees, especially for gynodioecious plants such as fig. In this study, genome resources were developed for *F. erecta* to facilitate fig breeding programs. The genome sequence of *F. erecta* was determined using single-molecule real-time sequencing technology. The resultant assembly spanned 331.6 Mb with 538 contigs and an N50 length of 1.9 Mb, from which 51,806 high-confidence genes were predicted. Pseudomolecule sequences corresponding to the chromosomes of *F. erecta* were established with a genetic map based on single nucleotide polymorphisms from double-digest restriction-site associated DNA sequencing. Subsequent linkage analysis and whole genome resequencing identified a candidate gene for the Ceratocystis canker resistance trait. Genome-wide genotyping analysis enabled selection of female lines that possessed resistance and effective elimination of donor genome from progeny. The genome resources provided in this study will accelerate and enhance disease resistance breeding programs in fig.

## Introduction

Common fig (*Ficus carica*: 2n = 2x = 26) has been cultivated for at least 11,000 years and is the oldest known cultivated plant (Kislev et al., 2006). It remains one of the most important fruit crops in the Mediterranean region due to its ability to thrive in dry climate conditions (Flaishman et al., 2008). However, fig is susceptible to devastating outbreaks of Ceratocystis canker, caused by the fungus *Ceratocystis ficicola*, when exposed to conditions of high temperature and humidity (Kajitani and Masuya, 2011). *C. ficicola* is thought to be soilborne (Kato et al., 1982). Fig saplings and young trees with Ceratocystis canker suffer wilting and poor growth of new branches, and infected trees eventually die (Morita et al., 2016; Shimizu and Miyoshi, 1999). Once Ceratocystis canker becomes established within an orchard, it becomes very challenging to eradicate due to the resilient chlamydospores produced by the fungus (Kato et al., 1982). One effective way to combat soil-transmitted diseases such as Ceratocystis canker is to develop resistant rootstocks. Highly resistant cultivars are therefore required for stable fig production in humid regions with high disease pressure such as Japan. However, while non-host resistance genes have been utilized in fig breeding programs (Himeno et al., 2015), no race-specific genes have been employed due to a lack of resistant genetic resources in common fig breeding materials.

A wild fig relative, *F. erecta* (2n = 2x = 26), possesses resistance to Ceratocystis canker (Morita et al., 2011; Shimizu and Miyoshi, 1999). However, introducing resistance characteristics into common fig from *F. erecta* has proved challenging. Hybrid breakdown is induced by interspecific crosses between common fig and *F. erecta*, and grafting compatibility between the two species is extremely low (Hosomi, 1993). Many attempts have been made to overcome these incompatibilities and generate interspecific hybrids between common fig and *F. erecta*. Yakushiji et al. (2012) succeeded in obtaining F1 hybrids that exhibited similarly high resistance to Ceratocystis canker as *F. erecta*; however, the plants had poor morphological phenotypes (Yakushiji et al., 2012). Recent research shows that Ceratocystis canker resistance in *F. erecta* is controlled by a single dominant gene (Yakushiji et al., 2019). The dominant nature of the resistance trait means that the F1 hybrids and progeny obtained in the earlier study could be used as rootstocks in common fig cultivation. Alternatively, repeated backcrossing would be expected to produce resistant fig cultivars with similar phenotypes to common figs. Due to the long generation time, production of new cultivars through repeated backcrossing is extremely time consuming. However, this can be expedited by background selection, in which residual chromosome segments derived from *F. erecta* are eliminated by genome-wide genotyping.

Fig is gynodioecious, having hermaphroditic caprifig-type trees (male flowers and short-style female flowers) and female fig-type trees (long-style female flowers) (Storey, 1975). Fig fruit crops are produced by the female fig-type trees. During breeding, therefore, selection of female plants as well as of those exhibiting favorable traits is desirable. Using whole genome sequence analysis, Mori et al. (2017) identified a candidate for the fig sex determinant gene, an ortholog of *RESPONSIVE-TO-ANTAGONIST1* (*RAN1*) in Arabidopsis (Hirayama et al., 1999), and developed a DNA marker linked to the locus, allowing identification of female seedlings prior to fruit bearing (Mori et al., 2017).

Genome resources are essential in breeding programs to develop fig cultivars with resistance to Ceratocystis canker, including for non-model/non-crop wild species such as *F. erecta*. Recent advances in sequencing technology have allowed genome sequences to be easily obtained for non-model and non-crop plants (Jiao and Schneeberger, 2017; Li and Harkess, 2018). In this study, the *F. erecta* genome sequence was obtained and characterized, providing a useful resource for fig breeding programs. Sequences were assigned to the chromosomes in accordance with a genetic map based on genome-wide single nucleotide polymorphisms obtained from a double-digest restriction-site associated DNA sequencing technique. A candidate gene for Ceratocystis canker was identified from an association analysis with chromosome-level sequences, and a DNA marker linked to the resistance locus were developed. The marker, alongside the previously identified sex determinant marker, can be used in a genome-wide genotyping technique and will allow rapid screening and development of Ceratocystis canker resistant fig cultivars.

## Results

### *Genome assembly and pseudomolecule sequence construction for* F. erecta

To estimate the size of the *F. erecta* genome, a kmer-distribution analysis was performed with short-read sequence data obtained from an *F. erecta* tree (Yakushiji et al., 2019; Yakushiji et al., 2012). The resultant distribution pattern indicated two peaks, representing homozygous (left peak) and heterozygous (right peak) genomes, respectively (Figure S1). The haploid genome of *F. erecta* was estimated to be 341.0 Mb in size. An initial genome assembly with a total of 21.3 Gb long reads (62.5× coverage of the estimated genome size) (Supplementary Table S1) consisted of 537 primary contig sequences and 1,917 haplotigs. The sequences were polished once to correct potential errors. The total length of the resultant primary contigs, designated as FER_r1.0pctg, was 331.6 Mb with an N50 of 1.9 Mb. The total length of the haplotigs, designated FER_r1.0hctg, was 264.2 Mb (N50 = 283.8 kb).

Assembly accuracy was validated with a genetic linkage analysis. To construct a genetic map, 24.6 Gb double-digest restriction-site associated DNA sequencing (ddRAD-Seq) reads were obtained for 121 B1F1 and parental plants (Supplementary Table S2). High-quality reads were aligned to FER_r1.0pctg as a reference with a map rate of 82.5% in average (Supplementary Table S2), and 13,320 SNPs were detected, which were then employed for linkage analysis. Thirteen linkage groups corresponding to the number of chromosomes of *F. erecta* were obtained, and marker order in each group and map distances between the markers were calculated. The resultant genetic map consisted of 12,748 SNP loci in 675 genetic bins covering 705.5 cM in total (Supplementary Table S3 and S4). In this mapping process, a probable missassembly point was found in contig Fer1.0_pctg0021F.1, whereby up- and downstream sequences were genetically mapped on two different linkage groups: 1a and 10. Therefore, this contig was broken into two sequences, Fer1.1_pctg0021F-1.1 and Fer1.1_pctg0021F-2.1, resulting in a final sequence total of 538. The sequence dataset was named FER_r1.1, and the primary contig and haplotig datasets were named FER_r1.1pctg and FER_r1.1hctg, respectively (Table 1). The total length of the resultant 538 primary contigs, FER_r1.1pctg, was 331.6 Mb with an N50 of 1.9 Mb. FER_r1.1pctg covered 97.2% of the estimated genome size. A Benchmarking Universal Single-Copy Orthologs (BUSCO) analysis indicated that 92.3% and 75.9% of complete BUSCOs were represented in FER_r1.1pctg and FER_r1.1hctg, respectively, reaching 94.6% when primary contigs and haplotigs were combined.

**Table 1.** Assembly statistics of the *F. erecta* genome.

|  | FER_r1.1 (Total) | FER_r1.1pctg | FER_r1.1hctg |
| --- | --- | --- | --- |
| Total scaffolds | 2,455 | 538 | 1,917 |
| Assembly size [bp] | 595,834,738 | 331,619,726 | 264,215,012 |
| Gaps size [bp] | 0 | 0 | 0 |
| Gaps % | 0 | 0 | 0 |
| N50 [bp] | 697,200 | 1,856,082 | 283,755 |
| N50 #sequences | 167 | 44 | 252 |
| N90 [bp] | 83,575 | 281,126 | 53,894 |
| N90 #sequences | 1,140 | 218 | 1,131 |
| Maximal scaffold [bp] | 9,725,511 | 9,725,511 | 1,827,087 |
| Complete BUSCOs | 94.6% | 92.3% | 75.9% |
| Single-copy | 33.2% | 87.0% | 70.6% |
| Duplicated | 61.4% | 5.3% | 5.3% |
| Fragmented BUSCOs | 1.3% | 2.2% | 2.8% |
| Missing BUSCOs | 4.1% | 5.5% | 21.3% |
| #Genes | 93,450 | 51,806 | 41,644 |
| Complete BUSCOs | 94.1% | 89.7% | 75.2% |
| Single-copy | 30.1% | 84.2% | 69.1% |
| Duplicated | 64.0% | 5.5% | 6.1% |
| Fragmented BUSCOs | 3.1% | 5.1% | 4.4% |
| Missing BUSCOs | 2.8% | 5.2% | 20.4% |

Based on the genetic map, 204 primary contig sequences spanning 273.2 Mb (82.4% of FER_r1.1pctg) were genetically anchored (Supplementary Table S5). The sequences were connected with ten thousand of Ns to construct pseudomolecule sequences, namely the FER_r1.1.pseudomolecule dataset. The names and directions of the pseudomolecule sequences were assigned in accordance with the previously described high-density genetic map for fig (Mori et al., 2017). Sequences that were unassigned to the genetic map were connected and termed chromosome 0 (FER_r1.1chr00).

### Prediction of protein-coding genes and repetitive sequences

In total, 80,323 and 61,755 protein-coding genes were initially predicted from the pseudomolecule sequences (FER_r1.1chr00–12) and haplotigs (FER_r1.1hctg), respectively. High-confidence genes were selected with two criteria: annotation edit distance score (Eilbeck et al., 2009) of ≤0.5 and amino-acid sequence length of ≥50. The filtered datasets included 51,806 and 41,644 genes in the pseudomolecule sequences (FER_r1.1chr00–12) and haplotigs (FER_r1.1hctg), respectively (Table 1). The gene set included 94.1% complete BUSCOs (89.7% and 75.2% for genes in the pseudomolecule sequences and haplotigs, respectively).

A predicted gene on chromosome 1a, Fer_r1.1chr01a_g003220.1, and a predicted haplotig gene, Fer1.1hctg_g318940.1, were counterparts of the candidate fig sex determinant gene s00259g14131.t1 (*RAN1*) (Mori et al., 2017). The proteins encoded by the two genes had histidine and glutamic acid at corresponding positions to amino acids 278 and 724 in s00259g14131.t1, respectively, both of which were encoded by the male allele in fig (Mori et al., 2017).

Repetitive sequences comprised 140.1 Mb (41.6%) and 108.0 Mb (38.1%) of the pseudomolecule sequences (FER_r1.1chr00–12) and haplotigs (FER_r1.1hctg), respectively (Supplementary Table S6). The dominant types in pseudomolecule sequences were LTR retroelements (48.2 Mb) followed by DNA transposons (23.4 Mb). Repeats sequences that were unavailable in public databases totaled 49.1 Mb.

### *Estimation of divergence time for* F. erecta

The 93,450 predicted genes in the *F. erecta* genome were clustered with those of fig (Mori et al., 2017), mulberry (He et al., 2013), jujube (Liu et al., 2014), and sweet cherry (Shirasawa et al., 2017) to obtain 23,102 clusters. Of these, 11,617 clusters were common across the five tested genomes, and 2,188 clusters consisting of one gene from each genome were selected for divergence time estimation. When the divergence time between fig and mulberry was set to 65 MYA (Kumar et al., 2017), the divergence time of *F. erecta* from the fig clade was 14.0 MYA (Supplementary Figure S2).

### *Sequence and structure variations in the* Ficus *genomes*

Sequence variations between the two haplotype sequences of the highly heterozygous *F. erecta* genome were detected by mapping haplotigs (FER_r1.0hctg) to the primary contigs (FER_r1.0pctg). In total, 1,893 haplotigs were mapped onto the 326 primary contigs and 1,400,071 variants were detected (Supplementary Table S7). Most of the variants (1,050,104; 75.0%) were in intergenic sequences, but 349,967 (25.0%) were found in gene regions. This included 31,867 deleterious variants (2.3%), such as frame-shift and non-sense mutations, in 11,278 genes.

Genome structure variations at an interspecies level were also investigated with respect to the fig genome (Mori et al., 2017). In total, 24,792 of the 27,995 contigs of the *F. carica* genome sequence were mapped onto the *F. erecta* pseudomolecule sequences, covering 193,530,279 bp in total (Supplementary Figure S3). Between the two genome sequences, 7,559,108 sequence variants were identified (Supplementary Table S7), of which 5,477,434 (72.5%) and 2,081,674 (27.5%) variants were located in intergenic and genic regions, respectively. Deleterious variants were found in 13,025 genes.

### Genetic mapping of the Ceratocystis canker resistance trait

Genotyping data from the BC1F1 plants used for genetic mapping were used for association analysis to find candidate genetic loci for Ceratocystis canker resistance. BC1F1 plants segregated into 62 resistant and 58 susceptible plants, fitting a 1:1 ratio (*p* = 0.715, chisq = 0.133) and indicating that the resistance trait was controlled by a single dominant gene (Yakushiji et al., 2019). As expected, a single significant peak (*p* = 2.03 × 10^-87^) was detected at the SNP at position 7,433,349 of chromosome 2 (Figure 1A–B). Furthermore, the *F. erecta* allele of the SNP (C) had a dominant effect for resistance over the fig allele (T).

**Figure 1.**
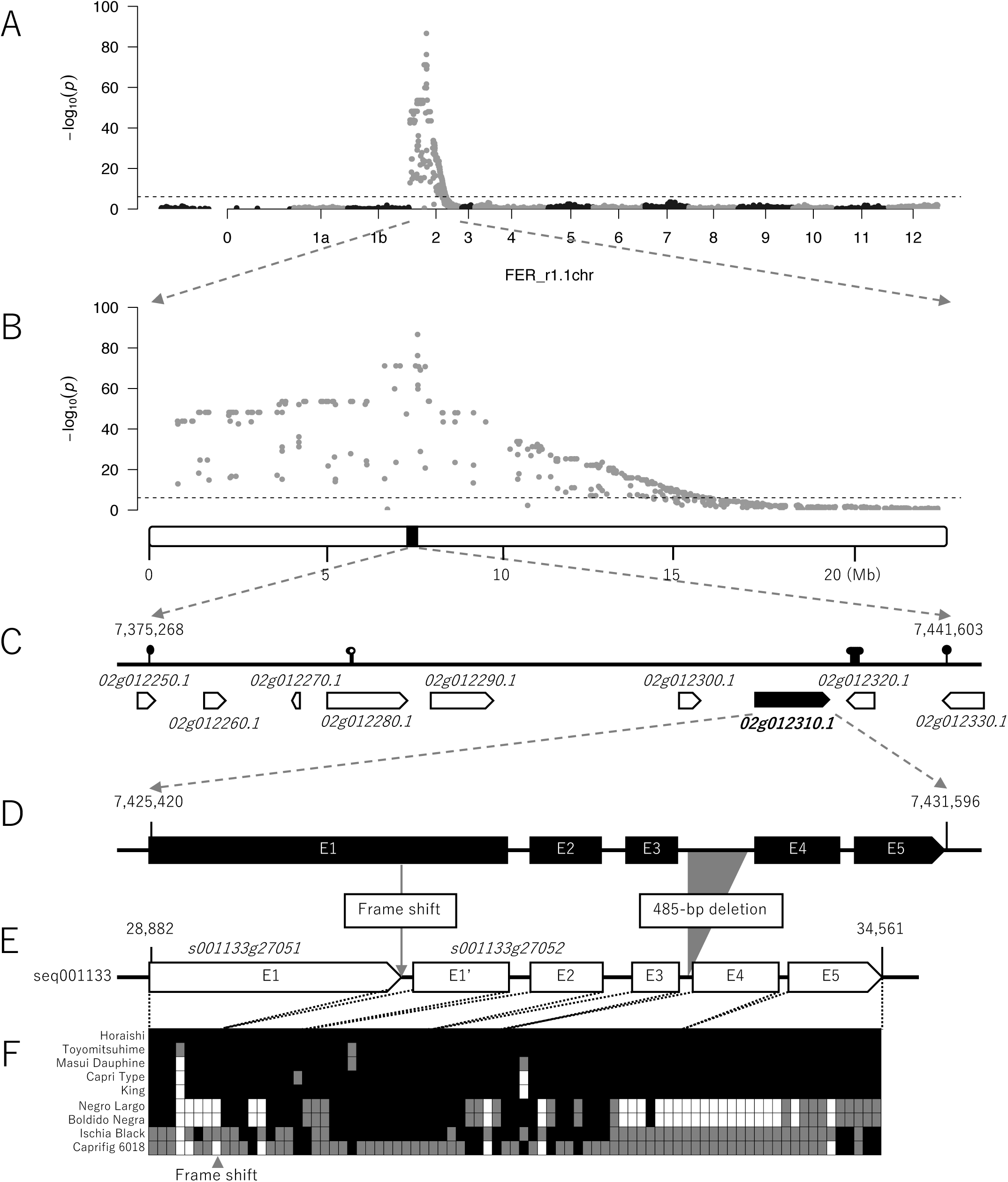
The Ceratocystis canker resistance locus in the *F. erecta* genome. **A and B.** Manhattan plots of genome-wide association for Ceratocystis canker resistance with resolution at the whole genome **(A)** and chromosome **(B)** levels. **C.** Gene order and direction in the candidate region for Ceratocystis canker resistance. Pins are SNPs detected by ddRAD-Seq. **D and E.** Gene structures of 02g012310.1 in the *F. erecta* genome **(D)** and s001133g27051 and s001133g27052 in the common fig genome **(E)**. Boxes indicate exons with exon numbers (E1 to E5), and major sequence variations are indicated. **F.** Genotypes of missense and frame-shift mutations among nine fig cultivars. Black and white indicate homozygotes of ‘Horaishi’ reference alleles and alternative alleles, respectively, while gray indicates heterozygotes. The frame-shift mutation disrupting 02g012310.1 is indicated with a triangle.

The chromosome recombination pattern of the BC1F1 population indicated that a 66.3 kb interval between positions 7,375,268 and 7,441,603 was the candidate for the resistance locus (Figure 1C). Nine genes were predicted in this region, one of which (Fer_r1.1chr02_g012310.1, 7,425,420–7,431,596) had five exons and was annotated as a nucleotide-binding adaptor shared by an apoptotic protease-activating factor-1 domain-containing disease-resistance protein (Figure 1D). The fig ‘Horaishi’ genome contained two counterpart genes, s01133g27051 and s01133g27052, from the 28,882–34561 region of the seq001133 scaffold sequence (Figure 1E). Between the two genome sequences, 198 SNPs and 11 indels (maximum size 485 bp at the third intron) were found. One of these, a single base deletion at position 7,427,300 of Fer_r1.1chr02 (position 30,762 of seq001133) caused a frame-shift mutation that split the single gene (Fer_r1.1chr02_g012280.1) in the *F. erecta* genome into two separate open reading frames (s01133g27051 and s01133g27052) in the fig ‘Horaishi’ genome. Furthermore, 40 and 72 missense mutations were found in s01133g27051 and s01133g27052, respectively, as interspecific sequence variations to Fer_r1.1chr02_g012310.1.

To investigate genotype patterns in fig cultivars, whole genome sequence data from five fig lines (‘Masui Dauphine’, ‘Negro Largo’, ‘Boldido Negra’, ‘Ischia Black’, and ‘Horaishi’) generated in this study and four lines (‘Caprifig 6018’, ‘Capri Type’, ‘King’, and ‘Toyomitsuhime’) generated previously (Mori et al., 2017) were mapped onto the fig ‘Horaishi’ genome sequence. Within the candidate gene region, 131 intraspecific variants were identified, but, unexpectedly, the frame-shift deletion was not conserved across the tested fig lines. Nevertheless, these genes still contained 78 missense mutations, one frame-shift mutation at the 3D end of s01133g27052, and one in-frame deletion, comprising nine different haplotypes in the nine tested lines (Figure 1F).

### Ceratocystis canker resistance breeding program

The single base deletion identified between *F. erecta* and ‘Horaishi’ fig was not conserved among the tested fig lines, and this mutation was therefore not useful for marker-assisted selection in fig breeding programs. Whole genome sequencing data revealed another polymorphism, at position 7,416,805 of FER_r1.1chr02, which was completely linked to the resistance phenotype. At this locus, resistant *F. erecta* was homozygous for ‘C’, whereas all nine susceptible fig lines were homozygous for ‘T’ as part of a recognition site for restriction enzyme BspHI, allowing a cleaved amplified polymorphic sequence (CAPS) marker to be developed. DNA fragments, including this SNP, were amplified by PCR and digested with BspHI. As expected, one DNA fragment was observed in *F. erecta*, whereas two and three fragments were detected in figs and in the F1 hybrid (FEBN-7), respectively (Supplementary Figure S4). This polymorphism was conserved in all accessions in a fig collection comprised of 122 lines (Mori et al., 2017). In addition, the CAPS genotyping scores in BC1F1 and BC2F1 plants completely matched the phenotypes, confirming the fitness of the marker for screening purposes.

Genome-wide genotypes of the BC1F1 and BC2F1 populations were investigated with ddRAD-Seq (Supplementary Table S2). As expected, the proportions of the genomes contributed by *F. erecta* and *F. carica* gradually decreased and increased, respectively (Supplementary Figure S5), with each generation from F1 to BC2F1 via BC1F1. Heterozygous genotypes also decreased. However, the genome proportions contributed by *F. erecta* and *F. carica* varied among the chromosomes (Supplementary Figure S6). In the BC2F1 population, full chromosomes 1b, 10, 11, and 12 and parts of chromosomes 2, 4, 5, 6, 7, 10, 11, and 12 were fixed with the *F. carica* genotypes; however, chromosome 3 retained a high proportion of the *F. erecta* genotype.

## Discussion

This study reports the first genome sequence for *F. erecta* and the second genome sequence for the *Ficus* genus following that of common fig, *F. carica* (Mori et al., 2017). Although the *F. erecta* genome is highly heterogeneous (Supplementary Figure S1), long-read sequencing allowed the *F. erecta* genome sequence to be assembled with high contiguity (Table 1). This contrasts with the common fig genome (Mori et al., 2017), which was highly fragmented (27,995 scaffold sequences with an N50 length of 166 kb) and covered only 70% of the estimated genome. Furthermore, chromosome-level pseudomolecule sequences were assembled for *F. erecta* in accordance with a genetic map based on SNPs from ddRAD-Seq analysis. The pseudomolecule sequences allowed association mapping to identify the genome position of the Ceratocystis canker resistance locus (Figure 1A–B).

The most likely candidate for the Ceratocystis canker resistance gene, Fer_r1.1chr02_g012310.1, was identified from the 66.3 kb resistance locus (Figure 1C). Fer_r1.1chr02_g012310.1 was selected because it was the only gene within the locus that was predicted to encode a disease resistance protein (Table 2). The counterpart in the fig ‘Horaishi’ genome was split into two genes, s01133g27051 and s01133g27052, as a result of a single base deletion that generated a frame-shift mutation leading to a premature stop codon (Figure 1D–E). We hypothesized that this split could be responsible for the differential disease susceptibility between the two species, but this variant was not observed across the nine fig cultivars tested (Figure 1F). Nevertheless, several mutations were identified in this gene in the other fig cultivars, namely 78 missense mutations, one frame-shift mutation, and one in-frame mutation, suggesting that multiple susceptible alleles may be present in common fig. This is consistent with observations that hyper-polymorphism is characteristic of resistance genes in plants (Ronald, 1998). The Fer_r1.1chr02_g012310.1 gene in *F. erecta* therefore remains a strong candidate for the Ceratocystis canker resistance gene. Further transcriptome analysis, transformation tests, and genetic approaches, such as high-resolution analysis with more diverse samples, would be required to validate this hypothesis.

**Table 2.** Gene annotation in the candidate genome region.

| Gene ID | UniProtKB<br>accession<br>number | Protein name | Organism name | E-value | High<br>impact<br>SNPs | Low<br>impact<br>SNPs | Moderate<br>impact<br>SNPs | Modifier<br>impact<br>SNPs |
| --- | --- | --- | --- | --- | --- | --- | --- | --- |
| Fer_r1.1chr02_g012250.1 | Q8SPU7 | Neuronal acetylcholine receptor subunit alpha-5 | *Bos taurus* | 0.064 | 1 | 3 | 2 | 18 |
| Fer_r1.1chr02_g012260.1 | Q54DY9 | Probable mitochondrial chaperone BCS1-B | *Dictyostelium discoideum* | 7.00E-23 | 0 | 8 | 3 | 13 |
| Fer_r1.1chr02_g012270.1 | O42941 | Peptidylprolyl isomerase cyp7 | *Schizosaccharomyces pombe* | 1 | 0 | 0 | 0 | 8 |
| Fer_r1.1chr02_g012280.1 | Q38942 | Protein RAE1 | *Arabidopsis thaliana* | 0 | 0 | 11 | 3 | 55 |
| Fer_r1.1chr02_g012290.1 | Q93WX6 | Cysteine desulfurase 2, chloroplastic | *Arabidopsis thaliana* | 0 | 0 | 5 | 3 | 42 |
| Fer_r1.1chr02_g012300.1 | Q9JK82 | Exostosin-1 | *Cricetulus griseus* | 1.6 | 0 | 1 | 0 | 21 |
| Fer_r1.1chr02_g012310.1 | Q9T048 | Disease resistance protein At4g27190 | *Arabidopsis thaliana* | 2.00E-75 | 1 | 36 | 74 | 17 |
| Fer_r1.1chr02_g012320.1 | Q9C899 | Feruloyl CoA ortho-hydroxylase 2 | *Arabidopsis thaliana* | 6.00E-81 | 0 | 4 | 3 | 4 |
| Fer_r1.1chr02_g012330.1 | Q9LHN8 | Feruloyl CoA ortho-hydroxylase 1 | *Arabidopsis thaliana* | 1.00E-111 | 0 | 2 | 12 | 18 |

Counterparts of the sex determinant gene candidate in common fig (Mori et al., 2017), s00259g14131.t1 (*RAN1*), were also found in the *F. erecta* genome sequences, Fer_r1.1chr01a_g003220.1 and Fer1.1hctg_g318940.1. Comparison of amino acids at key positions indicated that both genes represented the male allele, consistent with the male *F. erecta* tree used for genome analysis and supporting our hypothesis that *RAN1* is the gene responsible for sex determination in fig (Mori et al., 2017). The sexes of the BC1F1 and BC2F1 plants have not yet been determined due to the long juvenile stages in some lines, and linkage analysis has not been completed at the time of writing. However, preliminary results suggest that the sex phenotypes of the populations completely match the *RAN1* genotype (data not shown). In general, it is thought that the sex of plants as well as animals is controlled by heteromorphic sex chromosomes, such as XY or ZW systems, or by sex-linked genome regions in which recombination is highly suppressed at the kb- or Mb-scale (Charlesworth, 2016). It was also recently reported that a single gene or a single base mutation in homomorphic chromosomes is involved in sex determination in amphibia and *Seriola* fish (Koyama et al., 2019; Miura, 2017). It therefore remains possible that sex determination could be controlled by a single gene in some plants, one of which may be fig. Transformation tests as well as functional analysis would be required to confirm this possibility.

Genome-wide genotyping provided genome-scale graphical genotypes of fig breeding materials (Supplementary Figure S4). As expected, the proportions of the genomes contributed by the *F. erecta* donor decreased with each generation (Supplementary Figure S5). Theoretically, the donor genome proportions would average 50%, 25%, and 12.5% in the F1, BC1F1, and BC2F1 populations, respectively. Recurrent backcrossing is consequently time consuming, particularly in trees with long generation times. However, if outliers within a population could be selected with donor genome proportions that were unusually lower (or higher) than average, the donor genome could be eliminated from progeny more rapidly. Outlier selection could thereby accelerate recurrent backcrossing procedures even in trees. The availability of pseudomolecule sequences for fig will facilitate the use of this selection strategy in breeding programs for the development of common fig cultivars with Ceratocystis canker resistance.

The *F. erecta* genome characterized in this study provided insights into Ceratocystis canker resistance breeding strategies as well as identified responsible candidates for the resistance and sex determination genes. The genome resources will also be valuable for identifying the mechanisms underlying *F. erecta* resistance to other diseases or to pests such as nematodes (Hosomi, 1993) and may also contribute to our understanding of genome coevolution between *F. erecta* and the fig wasp (*Blastophaga nipponica*) (Wachi et al., 2016).

## Materials and Methods

### Plant materials

A male *F. erecta* tree was used for genome sequencing analysis. This tree had been used for an interspecific hybridization with *F. carica* ‘Boldido Negra’ to generate an interspecific F1 hybrid, i.e., FEBN-7 (Yakushiji et al., 2019; Yakushiji et al., 2012). The backcrossed lines (BC1F1, n = 121), derived from crosses between FEBN-7 and *F. carica* ‘Masui Dauphine’, ‘Negro Largo’, ‘Boldido Negra’, or ‘Ischia Black’ (Yakushiji et al., 2019), were used for linkage analysis (Supplementary Table S2). Advanced backcross lines (BC2F1, n = 114), generated by crossing a line of BC1F1 (MABN7-6) with either *F. carica* ‘Masui Dauphine’ or ‘Horaishi’ (Supplementary Table S2), were used for validation of the fitness of the DNA markers. Five fig cultivars, ‘Masui Dauphine’, ‘Negro Largo’, ‘Boldido Negra’, ‘Ischia Black’, and ‘Horaishi’, were used for whole genome resequencing analysis. A fig collection (n = 122) used in our previous study (Mori et al., 2017) was used to validate the resistance locus. The resistance levels of the accessions were evaluated as described previously (Yakushiji et al., 2019; Yakushiji et al., 2012).

### Genome sequencing analysis and assembly

High-molecular-weight genome DNA was extracted from young leaves of the *F. erecta* tree using the CTAB method (Murray and Thompson, 1980). Sequence libraries were prepared and sequenced using a Sequel system (PacBio, Menlo Park, CA, USA). The sequence reads were assembled by FALCON v. 1.8.8 (Chin et al., 2016) to generate primary contig sequences and associate contigs representing alternative alleles. Haplotype-resolved assemblies (i.e., haplotigs) were generated with FALCON_Unzip v. 1.8.8 (Chin et al., 2016). The resultant contig sequences were polished with ARROW v. 2.2.1 implemented in SMRT Link v5.0 (PacBio). Short-read data of *F. erecta* were used for genome size estimation with Jellyfish v. 2.1.4 (Marcais and Kingsford, 2011). Completeness of the assembly was assessed with sets of BUSCO v. 1.1b (Simao et al., 2015).

### Construction of genetic map-based pseudomolecule sequences

The BC1F1 population was analyzed with double-digest restriction-site associated DNA sequencing (Peterson et al., 2012). The library was prepared with PstI and MspI, as described previously (Mori et al., 2017), and sequenced using a HiSeq 2000 (Illumina, San Diego, CA, USA) system in paired-end, 93 bp mode. Data processing was also performed in accordance with Mori et al. (2017). High-quality reads were selected by trimming adapters with fastx_clipper (parameter, -a AGATCGGAAGAGC) in FASTX-Toolkit v. 0.0.13 (http://hannonlab.cshl.edu/fastx_toolkit) and deleting low-quality bases with PRINSEQ v. 0.20.4 (Schmieder and Edwards, 2011). Reads were aligned on FER_r1.1pctg with Bowtie2 v. 2.2.3 (Langmead and Salzberg, 2012), and sequence variants were detected with the mpileup command in SAMtools v. 0.1.19 (Li et al., 2009). High-confidence heterozygous SNPs in FEBN-7 were selected with VCFtools v. 0.1.12b (parameters of --minDP5 --minQ 999 --max-missing 0.75) (Danecek et al., 2011). A genetic map was constructed with LepMap3 v. 0.1 (Rastas, 2017). With the SNPs as anchors, the primary contig sequences were assigned to the linkage groups to establish the pseudomolecule sequences representing the chromosome sequences of *F. erecta*. The haplotigs as well as the contig sequences of the fig genome (Mori et al., 2017) were aligned on the pseudomolecule sequences with NUCmer of the MUMmer package v. 3.23 to detect sequence variations.

### Gene annotation and repeat detection

Total RNA was extracted from uninfected leaves and stems of *F. erecta* and from those artificially infected with *Ceratocystis ficicora*. Iso-Seq libraries were constructed and sequenced on the Sequel system (PacBio) according to the manufacturer’s protocol. The sequence reads were clustered to construct consensus sequences, also known as isoforms, with Iso-Seq2 pipeline in SMRT Link v. 5.1 (PacBio), and open reading frames (ORFs) were determined with ANGEL v. 2.3 (https://github.com/PacificBiosciences/ANGEL). Peptide sequences deduced from the ORFs and those predicted in the genomes of fig (Mori et al., 2017) and mulberry (He et al., 2013) were used for MAKER pipeline v. 2.31.10 (Cantarel et al., 2008). An AUGUSTUS training set from BUSCO analysis (Simao et al., 2015) with the *F. erecta* genome was also employed for the MAKER pipeline to predict putative protein-coding genes in the *F. erecta* genome. Repetitive sequences were detected with RepeatMasker v. 4.0.7 (http://www.repeatmasker.org), in which we used repeat sequences obtained from the *F. erecta* genome with RepeatModeler v. 1.0.11 (http://www.repeatmasker.org) and dataset registered in Repbase (Bao et al., 2015).

### Whole genome resequencing analysis

Genome DNA was extracted from young leaves of the *F. erecta* tree, three *F. carica* cultivars (‘Negro Largo’, ‘Boldido Negra’, and ‘Ischia Black’), and the F1 hybrid FEBN-7, using a DNeasy plant mini kit (Qiagen, Hilden, Germany). Genomic DNAs were used for library construction as described previously (Shirasawa et al., 2016). Sequences were obtained using a NextSeq 500 system in paired-end mode with a read length of 151 bp. In addition, previously generated whole genome sequence data from six lines (‘Caprifig 6018’, ‘Capri Type’, ‘Horaishi’, ‘King’, ‘Masui Dauphine’, and ‘Toyomitsuhime’) were also used (Mori et al., 2017). Data processing was performed as described for genetic mapping. High-quality reads obtained after trimming adapters and deleting low-quality bases were aligned on either the FER_r1.1. pseudomolecule or the fig genome contigs (GeneBank accession number: BDEM00000000) to detect high-confidence SNPs. The effects of mutations on gene function were predicted with SnpEff (v. 4.2; parameters: -no-downstream and -no-upstream) (Cingolani et al., 2012).

### Gene clustering, multiple sequence alignment, and divergent time estimation

Potential orthologs were identified from genes predicted in the *F. erecta* genome and from four genomes, fig (Mori et al., 2017), mulberry (He et al., 2013), jujube (Liu et al., 2014), and sweet cherry (Shirasawa et al., 2017), as an outgroup, using OrthoFinder v. 2.3.1 (Li et al., 2003). The single-copy orthologs in the five genomes were used to generate a multiple sequence alignment using MUSCLE v. 3.8.31 (Edgar, 2004), in which indels were eliminated by Gblocks v. 0.91b (Castresana, 2000). A maximum-likelihood algorithm-based phylogenetic tree was constructed from the alignments with the Jones-Taylor-Thornton model in MEGA X v. 10.0.5 (Kumar et al., 2018). The divergence time was calculated using MEGA X v. 10.0.5, assuming that the divergence time between fig and mulberry was approximately 65 MYA in TIMETREE (Kumar et al., 2017).

### Genome-wide association study

High-confidence SNPs used in the genetic mapping were used for genome-wide association analysis with GLM implemented in TASSEL 5 (Bradbury et al., 2007). Threshold values of *p*-value for association were adjusted with the Bonferroni correction.

### Genotyping analysis of the BC2F1 population

Genome-wide SNPs of the BC2F1 lines were genotyped with ddRAD-Seq as described above. A pair of oligonucleotide sequences (CGGCATCAGTTTCTTCATATTCT and CTGCACCGTTCTCTCTCTCC) was used as PCR primers for CAPS genotyping of SNPs at the resistance locus. CAPS analysis was performed as described previously (Mori et al., 2017).

## Data Availability

The sequence reads are available from the DNA Data Bank of Japan (DDBJ) Sequence Read Archive (DRA) under the accession numbers DRA008784, DRA008785, and DRA008786. The DDBJ accession numbers of assembled scaffold sequences are BKCH01000001-BKCH01002455. The genome information of *F. erecta* from this study and *F. carica* from our previous study (Mori et al., 2017) are available at Plant GARDEN (https://plantgarden.jp).

## Acknowledgments

We thank S. Sasamoto, S. Nakayama, A. Watanabe, T. Fujishiro, Y. Kishida, M. Kohara, C. Minami, H. Tsuruoka, and M. Yamada (Kazusa DNA Research Institute) for their technical assistance. This work was supported by the Kazusa DNA Research Institute Foundation and JSPS KAKENHI Grant Numbers 16H04878 and 16H06279.

