## Supplementary Figures for "The *Ficus erecta* genome to identify the Ceratocystis canker resistance gene for breeding programs in common fig (*F. carica*)"

**Supplementary Table S1.** Long reads for whole genome shotgun analysis in *F. erecta*
**Supplementary Table S2.** Short-read data for whole genome resequencing and ddRAD-Seq analysis
**Supplementary Table S3.** Genetic map of *F. erecta*
**Supplementary Table S4.** SNP loci on the genetic map
**Supplementary Table S5.** Summary of the *F. erecta* pseudomolecule sequence
**Supplementary Table S6.** Repetitive sequences in the *F. erecta* genome
**Supplementary Table S7.** Sequence variants in the *F. erecta* genome and impacts on gene functions

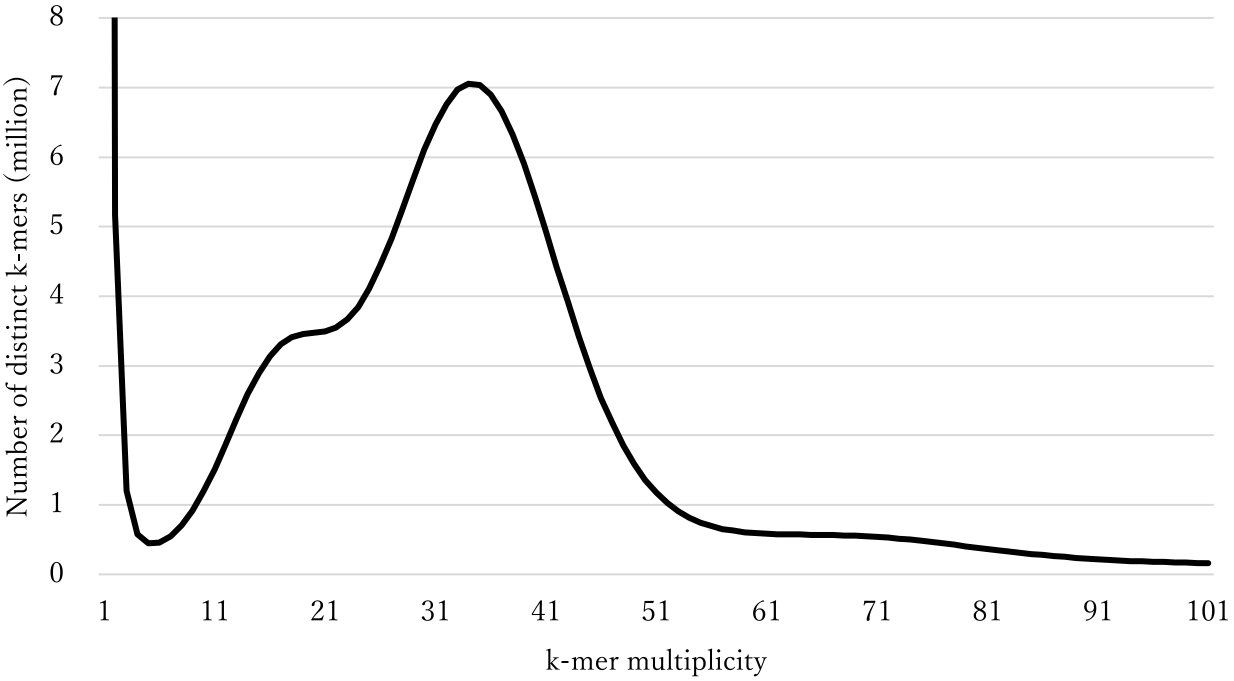


**Supplementary Figure S1.** Genome size estimation for *F. erecta* with the distribution of the number of distinct k-mers (k = 17) with the given multiplicity values.

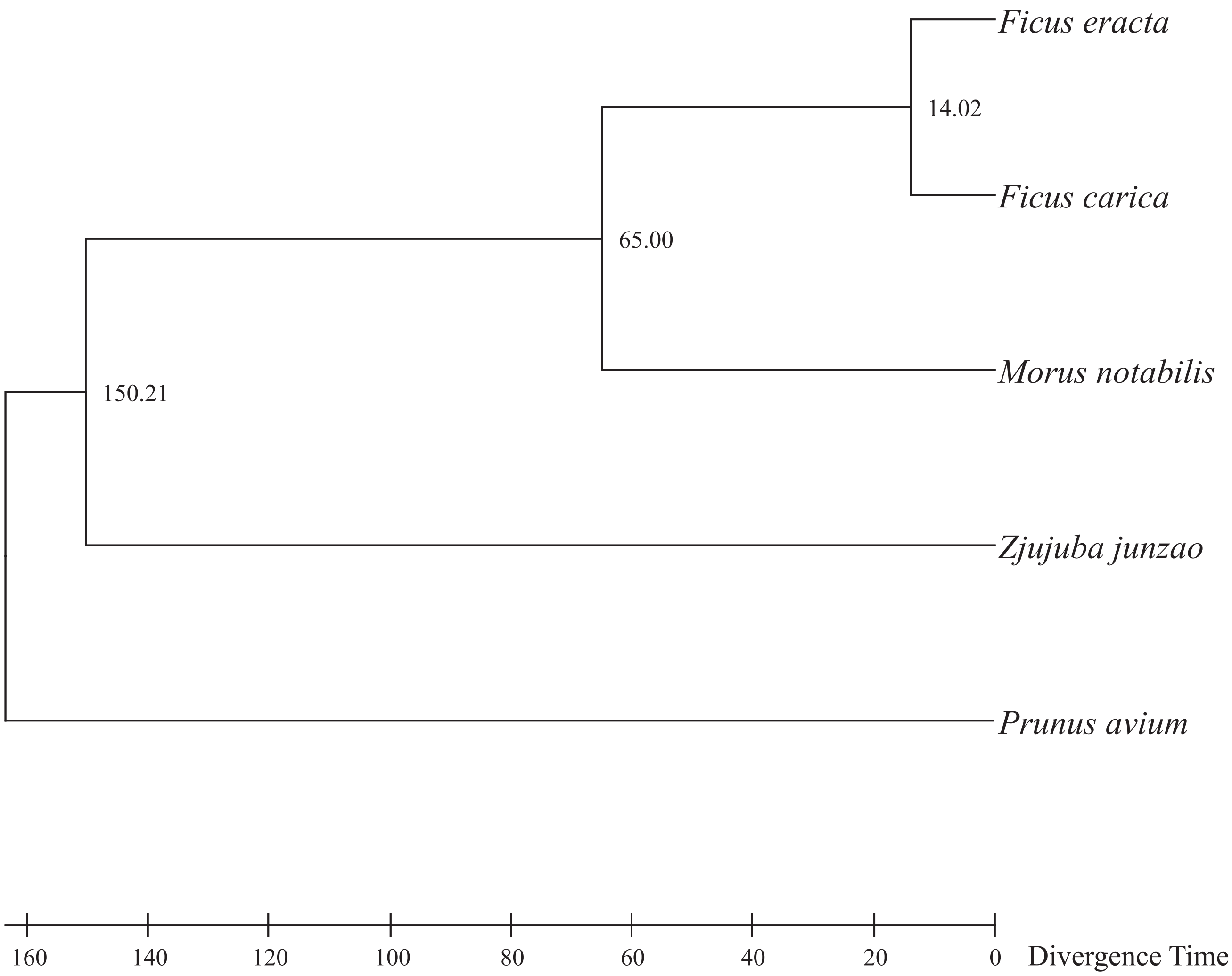


**Supplementary Figure S2.** Phylogenetic tree indicating the divergence time of *F. erecta.* Divergence times (MYA; million years ago) between branches are shown.

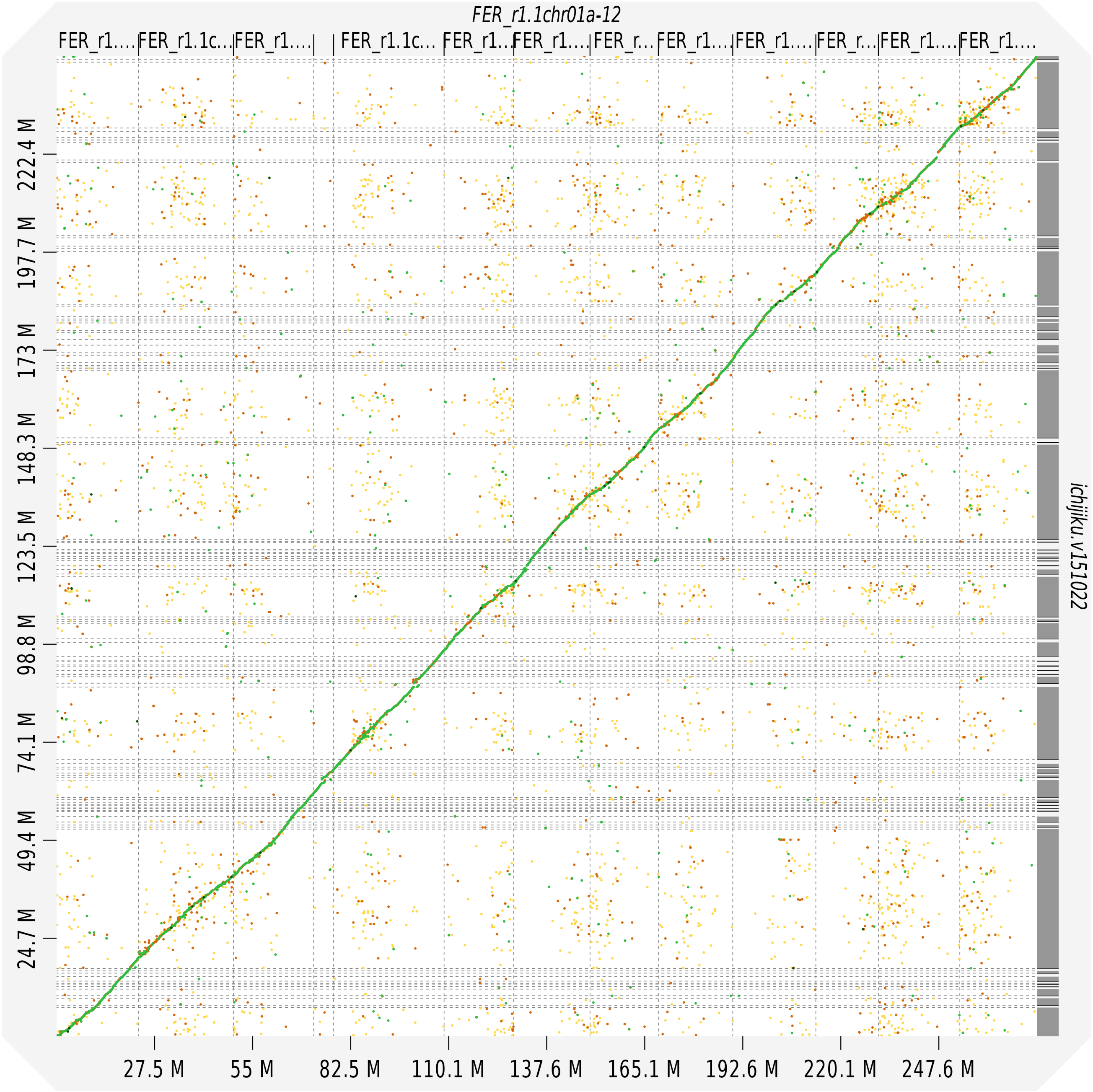


**Supplementary Figure S3.** Synteny of the genomes of *F. erecta* and common fig.

X-axis: the genome of *F. erecta* from this study; Y-axis: the genome of common fig (Mori et al. 2007).

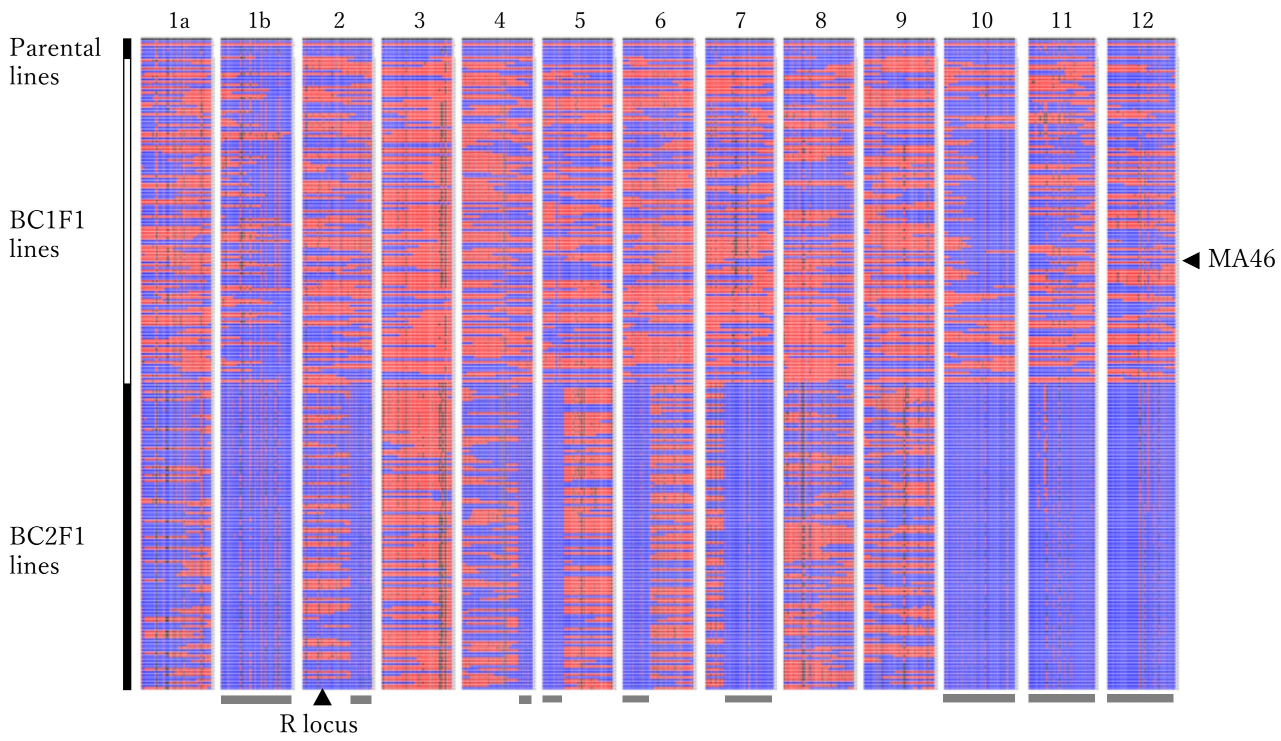


**Supplementary Figure S4.** Graphical genotypes of breeding materials.

Each row indicates breeding lines (parental lines, BC1F1 lines, and BC2F1 lines). The *F. erecta* chromosomes, 1a to 12, are shown as bars colored blue and red representing homozygotes of alleles of common fig and heterozygotes of *F. erecta* and common fig, respectively. The Ceratocystis canker resistance locus is indicated by a triangle. MA46 is the parental line of BC2F1. Gray bars (underneath) are chromosome regions fixed with the *F. carica* genotypes in BC2F1.

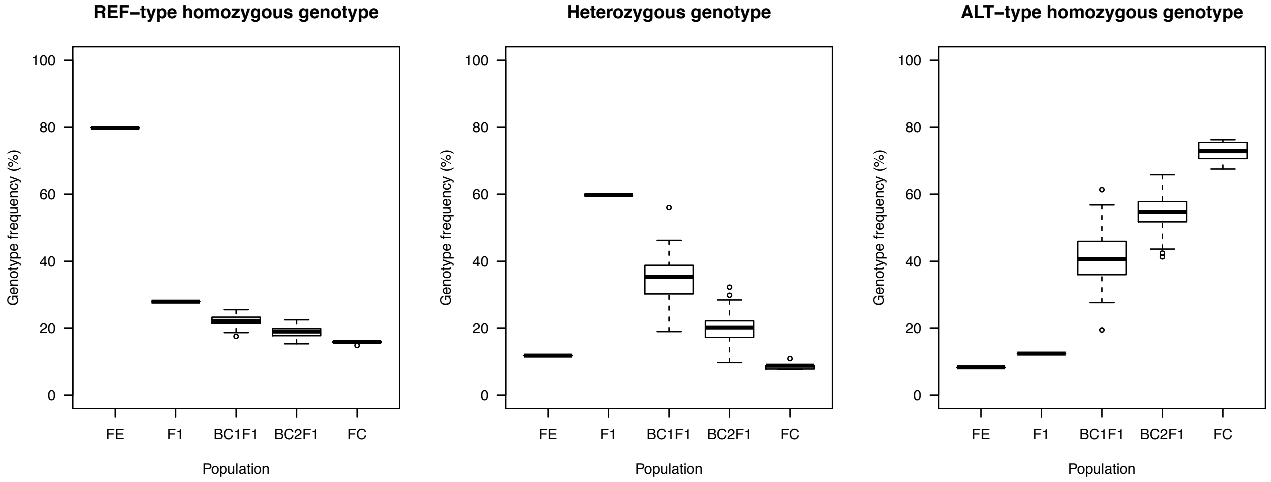


**Supplementary Figure S5.** Proportions of the genomes of *F. erecta* and common fig in the breeding populations.

FE, FC, and F1 indicate *F. erecta*, *F. carica* ('Boldido Negra', 'Horaishi', 'Ischia Black', 'Masui Dauphine', and 'Negro Largo'), and the F1 hybrid (FEBN-7).

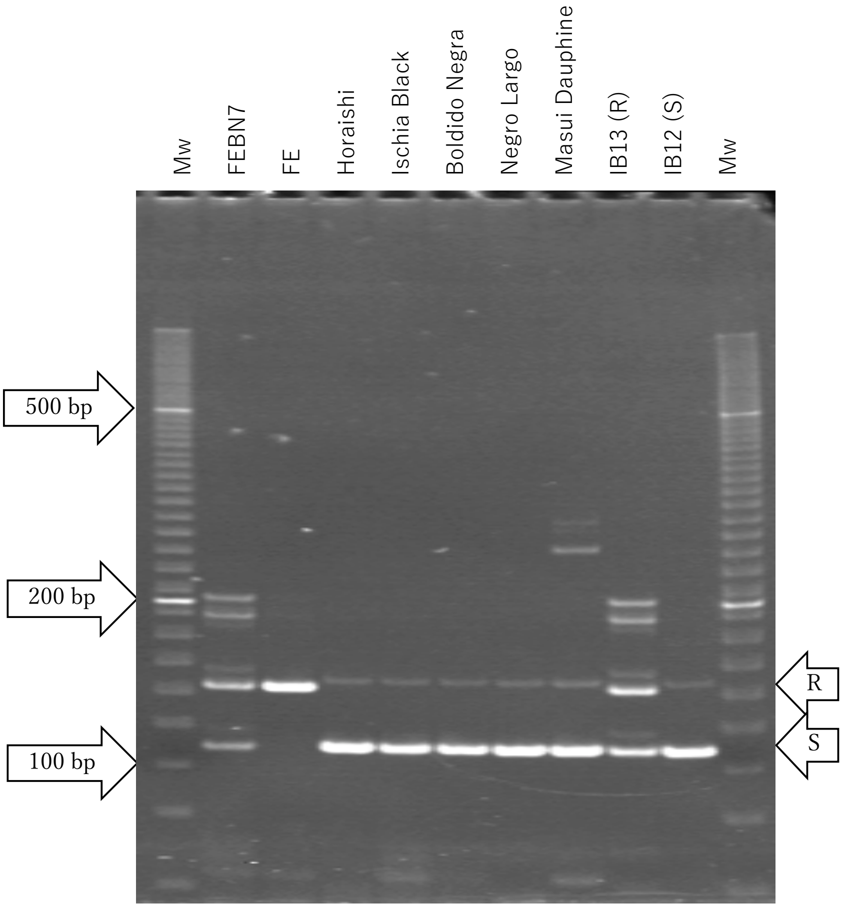


**Supplementary Figure S6.** The CAPS marker linked to Ceratocystis canker resistance
FEBN-7 is the F1 hybrid between *F. erecta* (FE) and *F. carica* ('Horaishi', 'Ischia Black', 'Boldido Negra', 'Negro Largo', and 'Masui Dauphine'). IB13 and IB12 are resistant and susceptible BC1F1 progeny. Mw indicates the molecular size marker, 20 bp ladder (Bio-Rad).
